# Skimming genomes for systematics and DNA barcodes of corals

**DOI:** 10.1101/2023.10.17.562770

**Authors:** Andrea M. Quattrini, Luke J. McCartin, Erin E. Easton, Jeremy Horowitz, Herman H. Wirshing, Hailey Bowers, Kenneth Mitchell, Makiri Sei, Catherine S. McFadden, Santiago Herrera

## Abstract

1: Numerous genomic methods developed over the past two decades have enabled the discovery and extraction of orthologous loci to help resolve phylogenetic relationships across various taxa and scales. Genome skimming (or low-coverage whole genome sequencing) remains a low-cost, promising method to not only extract high-copy loci, but also 100s to 1000s of phylogenetically informative single-copy nuclear loci (e.g., ultraconserved elements [UCEs] and exons) from contemporary and historical museum samples. The subphylum Anthozoa, which includes important ecosystem engineers (e.g., stony corals, black corals, anemones and octocorals) in the marine environment, is in critical need of phylogenetic resolution and thus might benefit from a genome-skimming approach.
2: Genome skimming was conducted on 242 hexacorals and octocorals collected from 1890 to 2022. Using previously developed target-capture baitsets, we bioinformatically obtained UCEs and exons from the genome-skimming data and incorporated them with data from previously published target-capture studies. We also extracted partial to whole mitogenomes and nuclear rRNA genes from the skim data.
3: The mean number of UCE and exon loci extracted from the genome skimming data was 1,837 ± 662 SD for octocorals and 1,422 ± 720 loci for hexacorals; phylogenetic relationships were well resolved within each class. A mean of 1,422 ± 720 loci were obtained from the historical museum specimens, with 1,253 loci recovered from the oldest specimen collected in 1886 and 1,336 loci recovered from a holotype. The nuclear *rRNA* genes and the majority of mitochondrial genes were successfully obtained from >95% of samples. Out of 99 circularized mitogenomes, 88% were recovered in samples from which we obtained >15M paired-end (PE) reads (>30M total reads); there was more variability in whether mitogenomes were circularized or not in samples with <15M PE reads.
4: Bioinformatically pulling UCEs, exons, mitochondrial genomes, and nuclear rRNA genes from genome skimming is a viable and low-cost option for phylogenetic studies. This approach can be used to review and support taxonomic revisions and reconstruct evolutionary histories, including historical museum and type specimens.

## Introduction

The advent of novel genomic methods and analyses has revolutionized our ability to resolve phylogenetic relationships across the tree of life. Numerous genomic methods (e.g., whole genome sequencing, transcriptomics, restriction-site associated sequencing, target-capture) developed over the past two decades have enabled the discovery and extraction of orthologous loci across multiple phyla. While high-quality whole genomes or transcriptomes are ideal in many situations, obtaining this genetic information from most animal taxa is still not technically feasible. But over the past decade, the average cost of high-throughput sequencing has rapidly decreased (Park & Kim, 2016). Now, we can multiplex many more taxa and obtain more genomic data (i.e., base pairs) per sample at a much lower cost than ever before. Therefore, genome skimming, or low-coverage whole genome sequencing (WGS), could be used to readily obtain enough orthologous loci, including conventional DNA barcodes, at a relatively low cost for phylogenomic studies (Trevisan et al., 2019; Liu et al., 2021).

Genome skimming has been used in prior studies to obtain whole mitochondrial genomes and nuclear DNA loci for phylogenetic studies (e.g., Malé et al., 2014; Liu et al., 2021; GoLightly et al., 2022; Taite et al., 2023). In addition, genome skimming has increasingly been used to help build DNA barcode reference databases for applications such as environmental DNA (eDNA) sequencing (Zeng et al., 2018; Hoban et al., 2022; Zhang et al., 2023). This method’s potential, however, for other applications remains unrealized, as typically more than 99% of the sequence data produced by skimming is not used (Bohmann et al., 2020). Low-coverage genome skims could readily be used to bioinformatically pull out ultraconserved elements (UCEs), exons, and other genes of interest. And because this method does not necessarily need high-quality DNA such as other methods (i.e., RAD Sequencing), genome skimming might be useful for historical samples that are housed in natural history museums across the globe (see Tin et al., 2014; Yeates et al., 2016; Bakker 2017; Liu et al., 2021; Hoban et al., 2022). Thus, this method should be more thoroughly explored for various applications across different qualities and quantities of genomic DNA.

Phylogenomic studies of marine invertebrates might benefit from a genome-skimming approach. In particular, the subphylum Anthozoa (*sensu* McFadden et al., 2022; phylum Cnidaria) is in critical need of taxonomic revision and resolution across family, genus, and species levels that will ultimately help in discriminating species and improving estimates of species diversity and distribution. Anthozoans are a diverse group of marine invertebrates, including sea anemones and corals, which are essential in building marine ecosystems from polar to tropical regions and the coasts to the abyss. Anthozoans currently comprise ∼7500 valid species (Daly et al., 2007) in two classes (Hexacorallia and Octocorallia, McFadden et al., 2022), but this number might be grossly underestimated (Plaisance et al., 2011; Bridge et al., 2023). Recently (i.e., in the past five years), the number of phylogenomic studies on anthozoans has grown rapidly. These studies have used a variety of methods, such as restriction-site associated sequencing (RADSeq, Reitzel et al., 2013; Herrera & Shank, 2016, Quattrini et al., 2019; Arrigoni et al., 2020), transcriptomics (Zapata et al., 2015), and target-capture genomics (e.g., Quattrini et al., 2020; Untiedt et al., 2019; Glon et al., 2021; McFadden et al., 2021, 2022; Bridge et al., 2023) to resolve questions at a range of scales. Target-capture of UCEs and exons, in particular, has shown much promise in resolving phylogenetic relationships of anthozoans across deep (i.e., orders, Quattrini et al., 2020; McFadden et al., 2021, 2022) to shallow (i.e., closely related species, Erickson et al., 2021; Glon et al., 2023; Bridge et al., 2023) time scales.

The original Anthozoa UCE and exons baitset was designed by Quattrini et al. (2018) and redesigned by Erickson et al. (2021) for Octocorallia and Cowman et al. (2021) for Hexacorallia. These baitsets target 1000s of loci, but do not include baits for mitochondrial genes or the nuclear ribosomal RNA (*rRNA*) genes. Although using mitochondrial genes and *rRNA* genes for phylogenomic studies of Anthozoa is cautioned (Figueroa & Baco 2015; Herrera & Shank, 2016; Quattrini et al., 2023), the utility of these markers goes beyond phylogenomic analyses. For example, mitogenome evolution across Anthozoa is intriguing as they exhibit a range of properties unique among metazoans, including gene order rearrangements (Brockman & McFadden, 2012; Lin et al., 2014; Figueroa & Baco, 2015; Seiblitz et al., 2022), a mismatch repair enzyme in Octocorallia (*mtMutS*, Bilewitch & Degnan, 2011), gene introns in the Hexacorallia (e.g., a homing endonuclease, Fukami et al., 2007, Barrett et al., 2020), and bipartite mitogenomes (Hogan et al., 2019). In some cases, mitogenomes have been used as taxonomic characters, as certain gene orders in mitogenomes appear to be restricted to particular families (see Seiblitz et al., 2022). Finally, with the recent efforts to monitor coral ecosystems with environmental DNA, there is a need to increase the number of taxa and loci in reference databases (McCartin et al., in review). Because genome skimming enables the production of low-coverage yet highly fragmented genomes, this method, followed by bioinformatic analyses, holds promise in not only obtaining whole mitogenomes and nuclear *rRNA* genes, but also UCEs and exons, as well as other genes of potential interest, from a range of DNA sample types (i.e., contemporary to historical samples) for a relatively low cost.

Here, we tested the utility of using genome-skimming data to bioinformatically obtain whole mitogenomes, nuclear *rRNA* genes, UCEs, and exons from hexacorals and octocorals. Although most of our efforts were focused on recently collected (< 20 years) specimens preserved specifically for genetic purposes, we also tested the utility of this approach to obtain UCEs, exons, and mitogenomes from historical material collected more than 100 years ago.

## Methods

### Collections

Octocorals (n=177) and hexacorals (n=32, including 30 antipatharians or black corals, one scleractinian [*Javania*], and one zoanthid [*Umimayanthus*]) were collected from the Gulf of Mexico, Caribbean Sea, and off the southeastern US coast from 2006 to 2019 on various expeditions. Specimens were collected with both Remotely Operated Vehicles (ROV) and SCUBA. Tissue samples were taken in the field, preserved in 95% ethanol and stored at -20°C, or flash frozen in liquid nitrogen and stored at -80°C. We also added historical, cataloged octocorals (n=33) collected from 1886 to 2006 from locations worldwide. Most museum specimens were preserved and/or stored dry or in 70% EtOH. See Supplemental Table 1 for further details.

### Molecular Lab Work

DNA was extracted in various ways (Table S1). Contemporary samples were extracted with either a modified CTAB protocol, a salting-out protocol, a GeneJet Genomic DNA Purification kit, or a Qiagen DNEasy extraction kit. Historical samples were all extracted with a Qiagen DNEasy kit. For some antipatharians and octocorals, DNA was cleaned with a Qiagen Power Clean Pro kit to remove PCR inhibitors. Samples were quantified with a fluorometer, either with a Quant-iT or Qubit.

For most samples (204 out of 242), library preparation was carried out in the Laboratories of Analytical Biology at the Smithsonian Institution. The quantity of genomic DNA input into a library preparation ranged from <0.65 ng to 93 ng total DNA; the average was 55±15 (SD) ng DNA. Library preparation was carried out using the NEBNext Ultra II FS DNA Library Prep Kit for inputs ≤ 100 ng with the following modifications: the reaction volume was reduced by half, the fragmentation/end prep incubation was conducted for 10 minutes (contemporary samples) or 2.5 minutes (historical samples), 5 μl of iTru Y-yoke adaptor (Glenn et al., 2019) was used instead of NEBNext Adaptor, adaptor ligation time was 30 minutes, bead cleanups were performed with KAPA Pure Beads, iTru i5 and i7 indices (Glenn et al., 2019) were used, and 10 cycles of PCR enrichment were conducted. A negative control was included on each plate during library preparation to test for any potential contamination. All DNA libraries were quantified and assessed with a Qubit fluorometer High Sensitivity Assay and a Tapestation, and final pools were created for sequencing on an Illumina NovaSeq (150 bp paired-end (PE) reads, Table S1). Pool 1 contained 33 historical samples sequenced on one lane of a NovaSeq S4 with 347 other invertebrate samples for a target read number of 5M PE reads per sample. Pool 2 contained 133 samples sequenced all together on one lane of a NovaSeq X for a target read number of 20M PE reads per sample. Pool 3 contained 38 samples sequenced with 57 additional samples on one lane of a NovaSeq X Plus for a target read number of 10M PE reads. To assess whether we could combine data from other DNA libraries, we included 38 DNA libraries (i.e., pool 4) that were prepared with an Illumina Nextera XT2 kit for NextSeq 500 sequencing at Biopolymers Facility at Harvard Medical School.

### UCE and Exon Analyses

Demultiplexed reads were trimmed using Trimmomatic v 0.32 or v 0.39 (Bolger et al., 2014). Trimmed reads were assembled using Spades v. 3.1 or 3.13.0 (Bankevich et al., 2012). Spades assemblies were then passed to phyluce v 1.7 (Faircloth 2016) to bioinformatically extract UCEs and exons using previously published bait sets for octocorals (octo-v2, Erickson et al., 2020) and hexacorals (hexa-v2, Cowman et al., 2020). The phyluce pipeline was used separately on octocorals and hexacorals as described in the online tutorials (https://phyluce.readthedocs.io/en/latest/tutorials/tutorial-1.html) with some modifications following Quattrini et al. (2018, 2020). Before aligning with MAFFT v7.130b (Katoh and Stanley 2013), we combined the data from 208 octocoral samples and the zoanthid *Umimayanthus* with previously published target-capture data obtained from 187 octocorals and 11 outgroups (Quattrini et al., 2018, 2020, Untiedt et al., 2020, Erickson et al., 2021, McFadden et al., 2022). We combined the data from 30 black coral samples and the stony coral *Javania* with previously published (Quattrini et al., 2018, 2020; Horowitz et al., 2022, 2023a) target-capture data from 106 black corals and four outgroups. After alignment, phyluce was used to create a 60% taxon-occupancy matrix for all loci, which were then concatenated separately for black coral (n=141) and octocoral (n=407) datasets. Phylogenomic analyses were conducted using maximum likelihood in IQTree v 2.1 (Nguyen et al., 2015) on the concatenated datasets with ultrafast bootstrapping (-bb 1000, Hoang et al., 2018) and the Sh-like approximate likelihood ratio test (-alrt 1000, SH-aLRT Anisimova et al., 2011). A partitioned model was used (-p). The best model of nucleotide substitution for each partition was found with ModelFinder (-m TESTMERGE, Kalyaanamoorthy et al., 2017) (Table 1). One octocoral sample, *Tripalea clavaria*, a dried museum specimen, was recovered as sister to all other octocorals. This sample was likely a contaminated sequence, which was pruned from the alignment. The alignment (n=406 species) was then re-run in IQTree using the abovementioned parameters.

### Mitogenome Analyses

For most samples (n=204), trimmed reads were also passed to Mitofinder v. 1.4 (Allio et al., 2020) for mitogenome assembly and annotation using a reference database of either octocorals or hexacorals downloaded from GenBank. We used trimmed reads in the analyses with the –new-genes parameter (to account for *mtMutS* and HEG) and the translation table 4 (-o). For the 38 samples from pool 4, mitogenomes were previously reported in Easton and Hicks (2019, 2020); thus, those results are not included in the present study.

### Nuclear rRNA Analyses

We also mapped, assembled, and extracted nuclear *rRNA* genes from the genome-skimming data. To obtain a reference sequence for mapping and assembly of octocoral samples, an annotated nuclear *rRNA* operon sequence, including the nuclear *rRNA* genes as well as *ITS1* and *ITS2*, was extracted from the NCBI-annotated *Xenia* sp. genome (RefSeq assembly GCF_021976095.1, scaffold NW_025813507.1) at NCBI (https://www.ncbi.nlm.nih.gov/genome/annotation_euk/all/). As a reference for black corals, we used a 4,721 bp sequence of *Cladopathes* cf. *plumosa* (GenBank: MT318868.1) from Barrett *et al*. (2020) that spans *18S*, *ITS1*, *5.8S*, *ITS2*, and the majority of *28S*.

Trimmed read pairs were merged using BBMerge v 38.84 (Bushnell et al., 2017) with the normal merge rate and the default settings and then imported into Geneious Prime v. 2023.1.2 (https://www.geneious.com). Merged read pairs were mapped and assembled to the reference sequences using the “Map to Reference(s)” function in Geneious with the sensitivity set to “Medium-Low Sensitivity/ Fast” and with five mapping iterations. Consensus sequences were generated from the resulting assemblies with the following settings. At each position, the threshold was set to 90% identity across all mapped reads for base-calling, a “?” was called if the coverage was less than 10 mapped reads, and the quality was assigned as the highest quality from any single base. Each consensus sequence was trimmed to its reference.

From the consensus sequences, we extracted and analyzed the *rRNA* genes *18S, 5.8S*, and *28S*. The consensus sequences were aligned using MAFFT v. 1.5.0 (algorithm E-INS-I, scoring matrix 100PAM/K=2) as implemented in Geneious Prime 2023.2.1 (https://www.geneious.com). Two alignments were analyzed, one including *ITS1* and *ITS2* in addition to the rRNA genes and another with *ITS1* and *ITS2* removed (e.g., *18S, 5.8S*, and *28S* only). The alignments were trimmed at the 5’ end to the beginning of *18S* using octocorals as a reference. While we were able to assemble the entirety of *28S* for octocorals, we were only able to assemble about one-half of the *28S* gene in black corals, due to incompleteness of the black coral reference sequence used. Partitions were created for both alignments (with and without the *ITS*). Phylogenetic inference was then conducted with IQTree using the best model of evolution determined by Modelfinder (-m TEST, Kalyaanamoorthy et al., 2017) and 1000 ultrafast bootstrap replicates (-bb 1000).

In addition to analyzing these concatenated *rRNA* gene alignments, we also extracted a ∼400 bp DNA barcode from the consensus sequences that is targeted by anthozoan-specific meta-barcoding primers (McCartin et al., in prep). This DNA barcode was compared to sequences generated via conventional PCR/Sanger sequencing for seven black coral and twenty-eight octocoral samples (McCartin et al., 2023). These barcoding sequences were aligned with MAFFT v. 7.49 (LINS-I method) and phylogenetic inference was conducted in the same manner using IQTree as for the concatenated alignment of *rRNA* gene sequences. Best models of sequence evolution for the partitioned datasets were chosen by ModelTest as implemented by IQTree (-m TEST)

### Statistical Tests

For historical museum specimens sequenced in pool 1, we conducted analyses to determine if collection year, library concentration, or DNA concentration impacted the number of reads or loci obtained. We first determined a significant correlation (r=0.58, p=0.001) between DNA and library concentration and thus removed DNA concentration from further analyses (Fig. 2A). Then, we assessed both additive and multiplicative linear regression models on log-transformed data to determine whether library concentration and collection year affected the dependent variables of number of reads and loci. The multiplicative models had a higher adjusted R-squared value (0.32, 0.69) than the additive models (0.24, 0.65) for tests on loci and read recovery, respectively; thus, we report the results of the multiplicative model below. We also tested whether the number of loci recovered was influenced by the number of reads obtained per sample.

We also determined whether the number of reads obtained across pools 1-3 significantly affected the completion of mitogenome circularization when using MitoFinder. We used a one-way analysis of variance on log-transformed data for both hexacorals and octocorals.

## Results

### Assembly Statistics

Of 242 samples, two failed sequencing with only 4,926 and 89,916 PE reads obtained; thus, these samples were removed from subsequent analyses. The remaining 240 samples had between 854,547 and 55,565,170 PE reads, with an average of 17,382,298 ± 8,065,341 PE reads. Pool 1 had an average of 8,343,203 ± 2,922,102; Pool 2 had an average of 23,156,985 ± 3,323,082; Pool 3 had an average of 12,822,312 ± 5,512,007; and Pool 4 had an average of 9,884,551 ± 857,465 PE reads. Trimmed reads were assembled into a mean of 741,347 ± 484,057 SD contigs per sample (range: 13,422 to 221,4702) (Table S1).

### UCE and Exon Results

UCEs and exons were successfully recovered from the genome skimming data of octocorals and hexacorals. For octocorals, 7 to 2,443 loci (mean 1,837 ± 662 SD) out of 3,023 targeted loci were recovered from each individual. The mean locus size was 1266 ± 1048 bp with a trend of increasing numbers of loci obtained with increasing numbers of PE reads until ∼10M PE reads, where the recovery rate reached a plateau (Fig. 1). Out of 206 octocorals, <200 loci were recovered in only 3% of samples; all of these samples were from pools 1 and 4 with a range of collection ages from 1960 to 2017 and a 10-fold range of obtained reads (973,960 to 9,534,512 PE reads).

**Figure 1.**
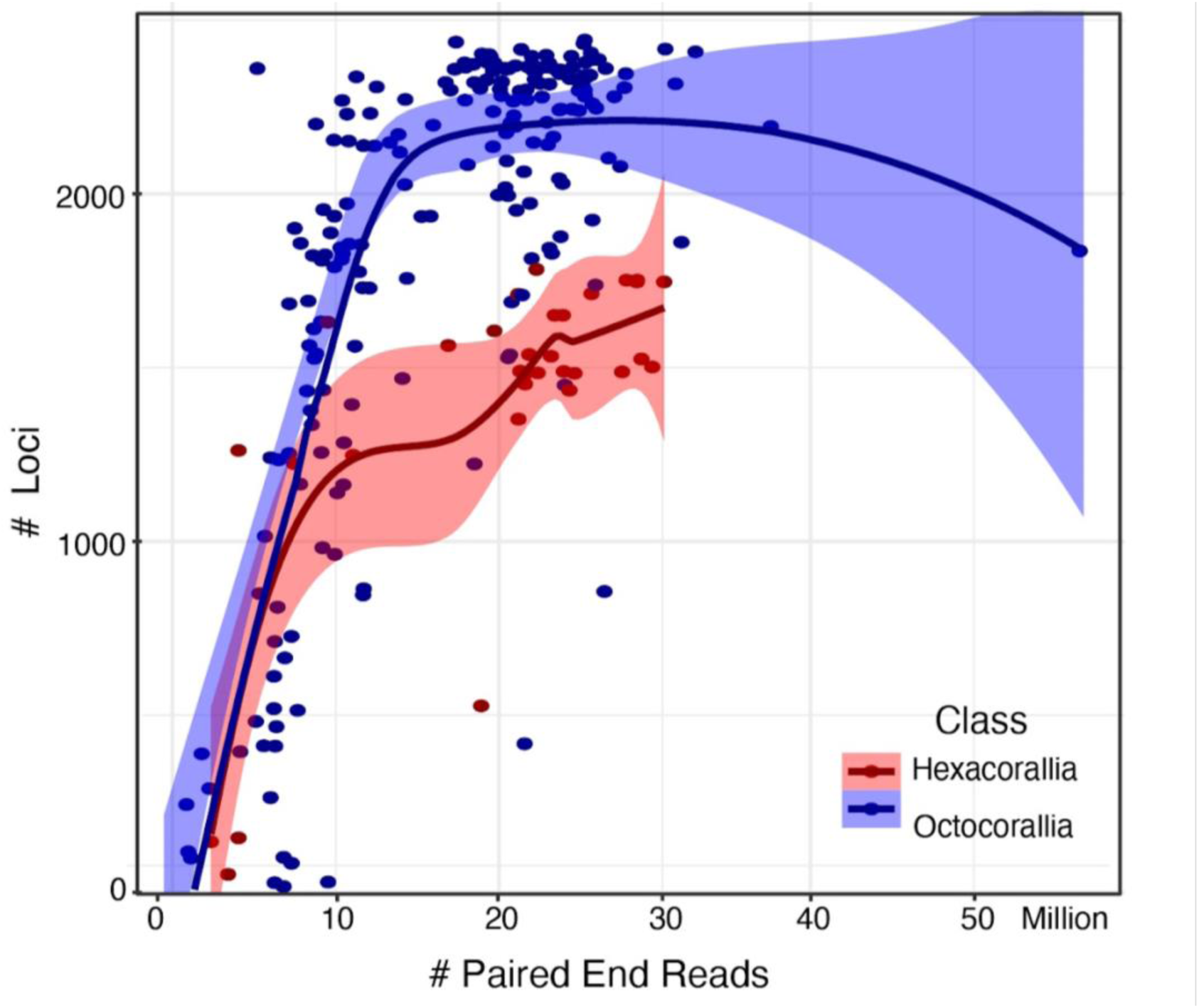
The number of loci recovered by the total number of paired-end reads obtained per sample in both Octocorallia and Hexacorallia. A polynomial regression model with local fitting was applied.

**Figure 2.**
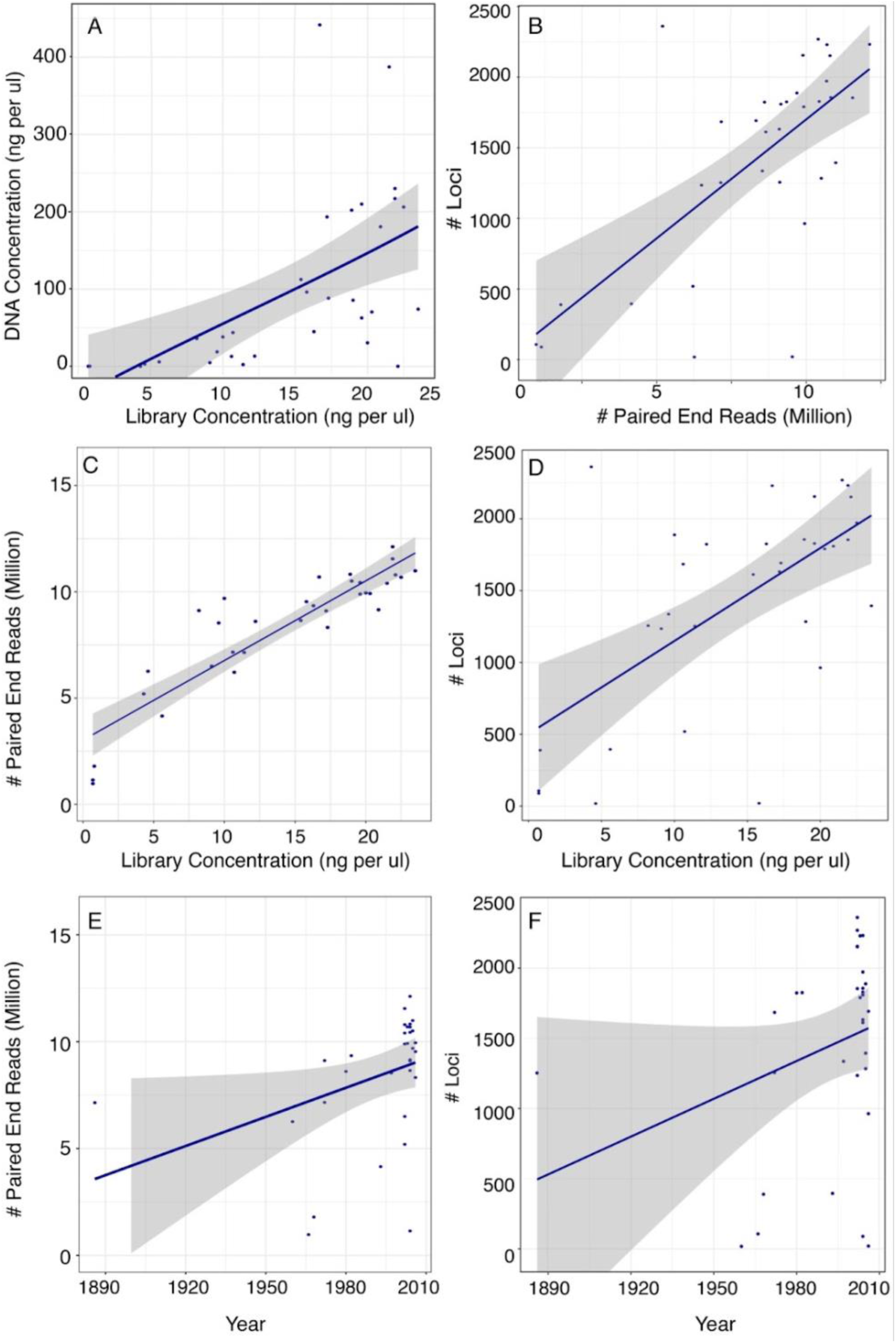
Data for historical museum samples sequenced in pool 1. (A) Library concentration versus DNA concentration. (B) Number of loci by the number of paired-end reads. (C-F) Number of reads and loci obtained by library concentration and collection year.

We were able to recover 18 to 2,361 loci (1,422 ± 720 loci) from the historical museum specimens, with 1,253 loci recovered from the oldest specimen collected in 1886 and 1,336 loci recovered from the holotype of *Sibogagorgia dennisgordoni* (Fig. 2). The mean locus size, however, was smaller (790 ± 578 bp) compared to the contemporary samples preserved specifically for genomics (1,355 ± 1,093 bp). In general, the number of loci recovered from the assemblies increased significantly (t=3.663, p=0.0009) with the number of reads obtained per specimen (Fig. 2B). The number of reads increased significantly with an increase in library concentration (t=2.31, p=0.028), with an interaction effect of year of collection (t=−2.25, p=0.032; Fig. 2C, E). Likewise, the number of loci increased significantly with an increase in library concentration (t=2.16, p=0.039), with an interaction effect of year of collection (t=-2.14, p=0.041; Fig. 2D, F)

The phylogenetic tree that included all octocoral samples from genome skimming and prior target-capture work (alignment: 1,262 loci, 243,326 bp) was well supported (Fig. 3, Fig. S1), and the genome-skimmed samples were recovered in the phylogeny within their respective families except one dried museum specimen, *Tripalea clavaria*, which was recovered as sister to all other octocorals and was thus pruned from the phylogeny. We recovered the two reciprocally-monophyletic orders, Scleralcyonacea and Malacalcyonacea, and added at least 55 species to the genomic-scale phylogeny of octocorals. Of 405 nodes, 96% had Sh-aLRT values over 80%, and 89% had bootstrap support values over 95%; most of the low values were near the tips. The zoantharian used as an outgroup in the octocoral phylogeny was correctly recovered in its respective order.

**Figure 3.**
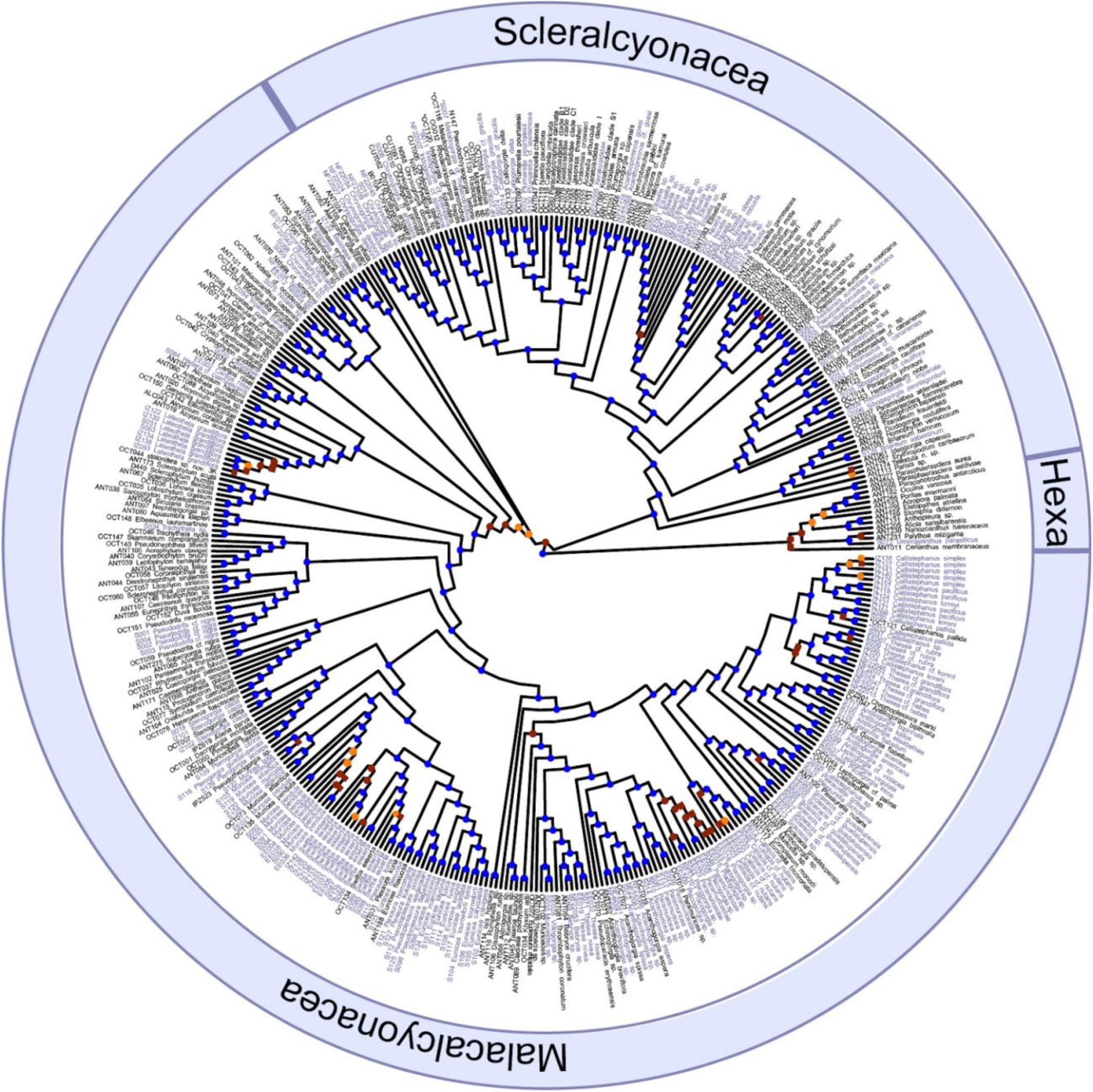
Maximum likelihood phylogeny of octocorals (purple=genome skim, black=target capture). Outgroups include hexacorals (Hexa). Node support values, represented by circles, include ultrafast bootstraps > 95% (blue), 80-95% (orange), and <80% (red). Where squares are indicated, SH-aLRT values were also < 80%. *=samples genome skimmed and target-enriched

For hexacorals, 42 to 1,783 loci (mean 1,379 ± 476 SD) out of 2,476 targeted loci were recovered from each individual. The mean locus size was 2,385 ± 1961 bp with a trend of increasing numbers of loci obtained with increasing numbers of PE reads until ∼20M PE reads, where the recovery rate slowed (Fig. 1). Out of 33 hexacorals, <200 loci were recovered in only 9% of samples; all of these samples were black corals collected in 2022 (sequenced in pool 2) with a range of obtained reads (2,353,550 to 4,045,520 PE reads).

The phylogenetic tree that included all antipatharian samples (alignment: 467 loci, 110,353 bp) from genome skimming and prior target-capture work was well supported, and the genome-skimmed samples were recovered in the phylogeny within their respective families (Fig. 4, Fig. S2). The newly incorporated genome-skim data (representing all seven antipatharian families) reinforces the monophyletic relationships of Myriopathidae and the monogeneric family, Leiopathidae. All other families are polyphyletic; notably, the new genome skim data reveals that Aphanipathidae is polyphyletic, where *Distichopathes hickersonae* and *Elatopathes abietina* are divergent from the rest of Aphanipathidae. This new dataset added at least 10 species to the black coral genomic-scale phylogeny. The scleractinian used as an outgroup in the hexacoral phylogeny was also recovered in its correct order. Out of 140 nodes, 70% had Sh-aLRT values over 80%, and 78% had bs values over 95%. In all cases, the lower node support values were near the tips.

**Figure 4.**
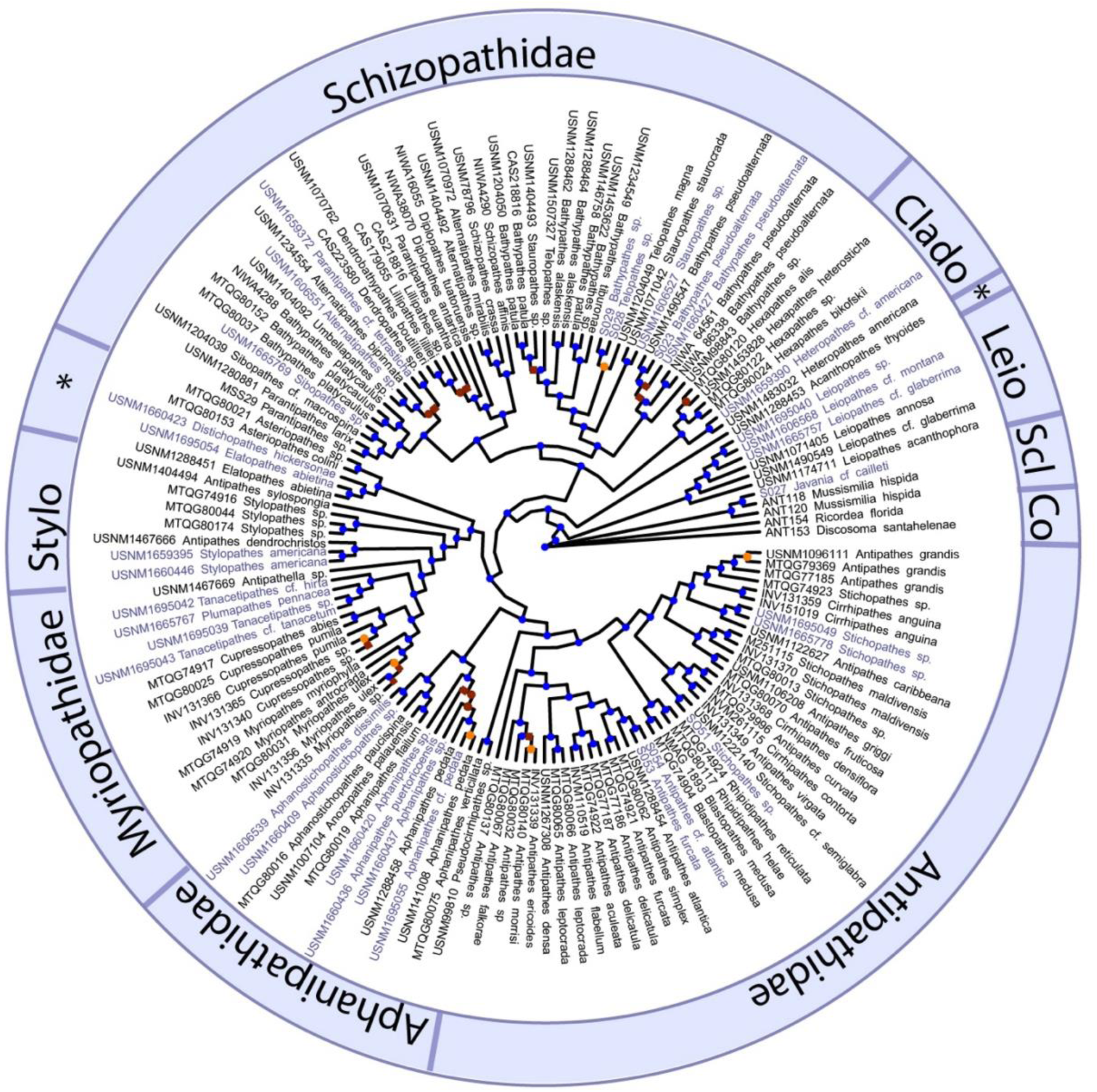
Maximum likelihood phylogeny of black corals (purple=genome skim, black=target capture). Outgroups include Scleractinia (Scl) and Corallimorpharia (Co). Node support values, represented by circles, include ultrafast bootstraps > 95% (blue), 80-95% (orange), and <80% (red). Where squares are indicated, SH-aLRT values were also < 80%. Leio=Leiopathidae, Clado=Cladopathidae, Stylo= Stylopathidae, and *= Species currently included within the polyphyletic family Aphanipathidae.

### Mitogenome Results

All mitochondrial protein-coding genes (PCGs) and *rRNA* genes were successfully retrieved from 95% of the samples targeted for mitogenome recovery. Of the 170 octocorals, we recovered 14 PCGs and both rSUs in 168 individuals. Only 10 PCGs and mitochondrial rSUs were recovered in two octocorals; both were museum samples collected in 1993 and 2005. The *mtMutS* sequences obtained were successfully integrated with data produced from PCR/Sanger sequencing, resulting in an alignment of 1074 bp (Fig. S3). Placements of taxa in the *mt MutS* phylogeny were as expected, and in many cases, the sequence data were 100% identical to the same species that were Sanger-sequenced. Most (70%) of octocoral mitogenomes were circularized with mitofinder. The majority of these were from Pool 2, which, on average, had the highest number of PE reads obtained across all pools (Fig. 5). Significantly more mitogenomes were circularized with a higher number of reads obtained for octocorals (ANOVA, F=96, p=0.001). For the 32 hexacorals, only one individual failed mitogenome assembly, with only three PCGs obtained, yet this individual had over 3,406,440 PE reads. Only 40% of all hexacoral mitogenomes were circularized with Mitofinder, with the majority of these from Pool 2. For hexacorals, no significant differences were found between mitogenome circularization and number of reads obtained (ANOVA, F=0.25, p>0.05).

**Figure 5.**
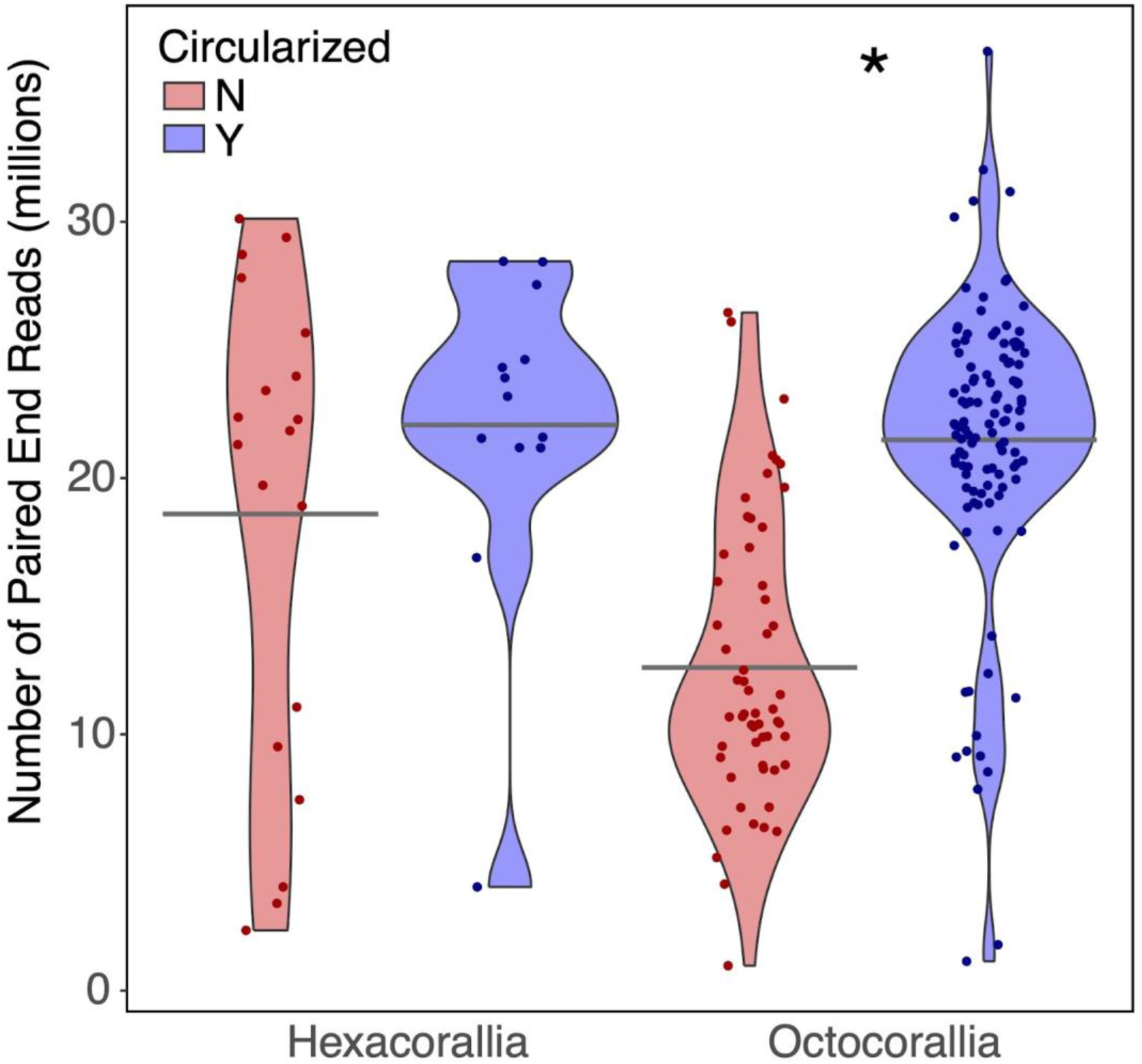
Circularization of mitogenomes by the number of paired-end reads from each sample for Octocorallia (n=171, 114 circular, 57 non-circular) and Hexacorallia (n=32, 13 circular, 19 non-circular). (*p=0.001). Gray bars indicate the group mean.

### Nuclear rRNA Results

Nuclear *rRNA* genes were successfully obtained from all but one sample. Reads mapped to at least 95% of the reference sequence used, and the mean coverage was 4,317x. The length of the assembled consensus sequences ranged from 4,142 to 6,136 bp, and differences in length were mainly because the black coral reference did not include all of the *28S*. Across the 478 bp alignment barcoding region of *28S*, sequences generated from genome skimming were 100% (p-distance) similar to their respective Sanger sequence except in the case of one specimen of *Sibogagorgia* cf. *cauliflora* (Fig. S4). This specimen had numerous ambiguous bases across the 478 bp alignment and was 15% divergent from the Sanger sequence. The phylogenetic tree produced from the *rRNA* genes (6,031 bp alignment) included taxa in positions as expected, except for *Sibogagorgia cf. caulifora* and *Plumarella pourtalesii (*Fig. S5).

## Discussion

### The Utility of Genome Skimming

Genome skimming is an effective approach for obtaining a range of loci useful for systematics and reference DNA barcode libraries of anthozoans. We bioinformatically obtained >1400 UCE/exon loci on average from both hexacorals and octocorals, indicating the utility of genome skimming for obtaining loci that are otherwise captured through a target-capture enrichment process. To highlight the similarity of results obtained from genome skimming and target capture, nine octocorals that were genome skimmed in this study were also target-captured in prior work (McFadden et al., 2022). In all cases, the numbers of UCE loci obtained from the skimmed samples were similar to or slightly higher (∼200 loci) than the target-captured samples, and five pairs of skimmed/target-captured samples included in the phylogeny were recovered as sister taxa. In addition to obtaining UCE/exon loci, we obtained nuclear ribosomal RNA gene sequences with high mapping coverage to reference sequences and most mitochondrial genes. Further, with minimal bioinformatic effort (i.e., just using one assembly program), we were able to obtain complete, circularized mitogenomes for 60% of all the samples. Our results indicate that this approach can also be used on historical-museum samples, where most target regions (i.e., mitochondrial genes, nuclear *rRNA* genes, UCES/exons) were successfully obtained.

Historical specimens, many of which had highly-degraded and low-quantity DNA (Table S1), performed very well with genome skimming. None of these historical specimens were preserved specifically for genetic applications. Yet, we recovered most, if not all, mt genes, nuclear *rRNA* genes, and more than 1000 UCE/exon loci from 75% of the samples. In addition, this approach is useful for obtaining numerous loci from type specimens (i.e., *Sibogagorgia dennisgordoni)* and specimens collected over 100 years ago. Our results, however, suggest that DNA concentration is directly correlated with library concentration, and higher library concentrations yield more reads and, thus, more UCE/exon loci. In contrast to expectations, the collection year had minimal impacts on UCE/exon loci obtained from the skimming data. Museum specimens used in this study were preserved in various ways, including drying, 70% EtOH, and 95% EtOH. Some specimens were likely fixed in formalin, but this information is often not retained in museum records. Thus, preservation type could confound a direct relationship of collection year with the number of loci obtained. Therefore, we recommend that researchers try genome skimming on various museum samples, regardless of collection age or preservation type. We also urge the use of type material in genome skimming studies to help resolve taxonomic issues in both classes of hexacorals and octocorals. Because the first step of preparing NEB genomic libraries is shearing DNA, one can skip or reduce the shearing time and use just the degraded DNA that is recovered from museum specimens in the DNA library preparation workflow. Our results here suggest that genome skimming is a simple genomic approach that can help unlock our historical museum collections, thus ultimately helping to resolve phylogenetic relationships across Metazoa.

There have been increasing efforts to use environmental (e)DNA sampling to characterize biodiversity and monitor health across ecosystems. It is clear, however, that the classification of eDNA sequences at a meaningful taxonomic resolution relies on the completeness of reference databases of DNA barcodes to which eDNA can be compared (Gold et al., 2021). But DNA barcodes remain missing for many metazoan taxa (e.g., Ransome et al., 2017; Pappalardo et al., 2021), and there are no standard barcodes that can be used to resolve species or even genera across diverse taxa, although both mitochondrial genes and nuclear *rRNA* genes are often used. Therefore, it is a critical time to increase barcode data across both taxa and gene regions. Our results suggest that genome skimming is one way to improve reference sequence databases simply and rapidly for applications like eDNA metabarcoding. We provide evidence that *28S rRNA* sequences recovered from the genome skimming data were largely congruent to sequences generated from conventional PCR amplification and Sanger sequencing. Another obvious advantage of genome skimming over Sanger sequencing is the ability to generate sequencing data for multiple barcoding genes simultaneously, including nuclear *rRNA* and mitochondrial genes.

For the amount of data obtained, genome skimming is a relatively cost-effective method compared to other genomic and genetic approaches. Library preparation, sequencing (10-20M PE reads), and quantification cost ∼$60-75 USD for this study. This same amount would facilitate sequencing ∼6-7 loci (approximate costs, $6-8 for sequencing, $5 for PCR reaction) through traditional PCR and Sanger. Although the average costs of genome skimming are relatively low compared to Sanger sequencing, the high costs and/or access to genomic sequencing facilities, high-performance computing, and bioinformatics training might still be prohibitive for some researchers, particularly those in low-income countries (see, e.g., Rana et al., 2020; Yek et al., 2022; Whiteford et al., 2023). By collaborating across international borders, we can easily pool samples from several research groups for sequencing at a genomic sequencing facility, at least in situations where DNA exchange restrictions are not an issue.

### New insights into octocoral phylogeny

At the genus level, the phylogeny of octocorals constructed here using a combination of data obtained from target-enrichment of conserved elements and genome skimming was largely congruent with that published previously using data from target-enrichment only (McFadden et al., 2022). Relationships among families were also mostly in agreement with that previous analysis, with the most notable exception being the recovery of the family Cladiellidae, sister to the gorgonian families Euplexauridae and Paramuriceidae, as was also found by Quattrini et al. (2023). The subordinal-level clades defined by McFadden et al. (2022) were not, however, as well supported by the analysis presented here (Fig. 3). These differences may be attributable to differences between analyses in taxon sampling or the numbers and identities of loci included (i.e., including saturated loci) and exemplify the challenges inherent in resolving the deepest nodes in a group of organisms that evolved in the pre-Cambrian (McFadden et al., 2021).

Genomic data were obtained for the first time from representatives of 11 genera (*Paracalyptrophora* Kinoshita, 1908; *Nicella* Gray, 1870; *Iciligorgia* Duchassaing, 1870; *Lateothela* Moore et al., 2017; *Hedera* Conti-Jerpe & Freshwater, 2017; *Chromoplexaura* Williams, 2013; *Pseudoplexaura* Wright & Studer, 1889; *Placogorgia* Wright & Studer, 1889; *Villogorgia* Duchassaing & Michelotti, 1860; *Aliena* Breedy et al., 2023; and *Thesea* Duchassaing & Michelotti, 1860). Phylogenetic placement of each of these genera was congruent with expectations based on previous phylogenetic analyses of mitochondrial and nuclear *rRNA* gene trees (Cairns & Wirshing 2018; McFadden et al., 2022; Breedy et al., 2023). The phylogenomic analysis recovered *Thesea* as polyphyletic, with some species grouping in the family Paramuriceidae and others in the Gorgoniidae, which is also congruent with previous phylogenetic analyses (Carpinelli et al., 2022). The paraphyletic relationships of *Gorgonia* to *Antillogorgia* and of *Plexaura* and *Pseudoplexaura* to *Eunicea* have also been recovered in previous studies (Grajales et al., 2007; Torres-Suarez 2014), as has the polyphyly exhibited by *Leptogorgia* (Poliseno et al., 2017).

Molecular data were obtained for the first time for four genera, allowing their familial relationships to be assessed. *Acanthoprimnoa* Cairns & Bayer, 2004, a genus whose membership in Primnoidae has never been questioned (Cairns & Bayer, 2004; Cairns & Wirshing, 2018), was instead found to be sister to Ifalukellidae. *Tripalea* Bayer, 1955, placed in Spongiodermidae based on morphology (Cairns & Wirshing 2015), appears instead to belong to Incrustatidae in the order Malacalcyonacea. Finally, *Caliacis* Deichmann, 1936 and *Pseudothelogorgia* van Ofwegen, 1991, genera whose familial affinities were left *incertae sedis* by McFadden et al. (2022), each occupy unique positions within the clade of malacalcyonacean gorgonians (clade 8 of McFadden et al., 2022), suggesting they each deserve family status. Before proposing those new families, however, it will be necessary to confirm the species-level identification of the material we sequenced by comparison to original type material.

### New insights into antipatharian phylogeny

The black coral phylogeny is mostly congruent with previous reconstructions (Horowitz et al., 2022, 2023b); however, this study includes three genera (*Distichopathes* Opresko, 2004, *Plumapathes* Opresko, 2001, and *Tanacetipathes* Opresko, 2001) that have been sequenced for the first time with high-throughput genomic techniques, providing new insights into phylogenomic relationships within the order. *Distichopathes* Opresko, 2004 was recovered sister to *Elatopathes* Opresko, 2004. Along with *Asteriopathes* Opresko, 2004, these three genera are currently placed in Aphanipathidae Opresko, 2004, but they form a monophyletic clade divergent from the rest of Aphanipathidae. Instead, the three genera show affinity to Stylopathidae Opresko, 2006, a finding consistent with Opresko et al. (2020) based on three mitochondrial and three nuclear gene regions. The recovered polyphyletic relationship of *Plumapathes* Opresko, 2001 and *Tanacetipathes* Opresko, 2001 (both of which reside in Myriopathidae Opresko, 2001) is notable as they possess distinctly different branching characteristics (planar in *Plumapathes* vs bottlebrush in *Tanacetipathes*). However, Horowitz et al. (2023b) emphasized that smaller-scale features, such as polyps and spines, are often more informative than branching characteristics. Most species within the Myriopathidae have very similar spine and polyp characteristics. Thus, these genera within Myriopathidae require further examination for a possible taxonomic revision.

Six out of the seven families in the order are polyphyletic based on this and previous phylogenetic reconstructions (Brugler et al. 2013, Horowitz et al. 2022, 2023a). Notably, the family Aphanipathidae contains genera spread across the tree (identified by ‘*’ in Fig. 4), highlighting the need for taxonomic revisions. However, a formal taxonomic review cannot be conducted because the type for Aphanipathidae by subsequent designation, *Aphanipathes sarothamnoides* Brook, 1889 has yet to be sequenced. Therefore it is not yet certain which clade represents the Aphanipathidae. This study demonstrates that genome skimming and target enrichment are suitable methods to yield high phylogenetic resolution of antipatharians. All that is needed now are sequence data from holotype or topotype material representing each nominal and currently accepted genus to fill gaps and better support taxonomic revisions.

## Acknowledgments

Gulf of Mexico collections were funded by the NOAA’s National Centers for Coastal Ocean Science, Competitive Research Program, the Office of Ocean Exploration and Research, and the RESTORE Science Program under awards NA18NOS4780166, NA18OAR0110289, and NA17NOS4510096 to PI S. Herrera, and by the Flower Garden Banks National Marine Sanctuary, Schmidt Ocean Institute, Texas SeaGrant, and Texas Parks and Wildlife awards to David Hicks (lead PI or co-PI). Southeastern US collections were funded by the DEEPSEARCH program (lead PI, E. Cordes), funded by the U.S. Department of the Interior, Bureau of Ocean Energy Management (BOEM), Environmental Studies Program, Washington, DC, under Contract Number M17PC00009. Puerto Rico specimen collections were funded by NOAA Ocean Exploration (PI, A. Quattrini), NOAA Fisheries, and the Smithsonian Women’s Committee. Specimens from Florida were collected during the 2019 Workshop on Caribbean Octocoral Biology funded by NSF OCE-1756381 to H. Lasker and P. Edmunds under Florida Fish & Wildlife Conservation Commission Special Activity License SAL18-2052A-SR. Collections in Panama were funded by the Cnidarian Tree of Life project, NSF EF-0531779 to P. Cartwright under the auspices of the Smithsonian Tropical Research Institute. Research activities in Puerto Rico waters were coordinated with the Department of Natural and Environmental Resources of Puerto Rico under permit #2022-IC-010. Research activities in the Flower Garden Banks National Marine Sanctuary were done per permits FGBNMS-2017-007-A2 and FGBNMS-2019-003-A2. Genomic sequencing was funded by NOAA OE, NOAA Office of Education Educational Partnership Program with Minority Serving Institutions awards NA16SEC4810009 and NA21SEC4810004, and BOEM. We thank S. Cairns, S. Rowley and R. Cordeiro for help with some species identifications.

## Data availability

Raw sequence reads were submitted to GenBank under BioProject #XXXXX, and individual genes can be found XXXX. All code can be found on GitHub https://github.com/quattrinia/GenomeSkim_paper, and alignment and tree files can be found on Figshare #10.25573/data.24319078

## Author contributions

AMQ, SH, and CSM conceived the study and designed the methodology. AMQ conducted genomic analyses, analyzed data, created figures, and wrote the manuscript. LJM, HB, JH, KM, and EEE conducted genomic analyses. SH, CSM, JH, and EEE provided data and taxonomic identifications. MS and HHW constructed DNA libraries. All authors edited and approved the final version.

